# Mad dephosphorylation at the nuclear envelope is essential for asymmetric stem cell division

**DOI:** 10.1101/798116

**Authors:** Justin Sardi, Muhammed Burak Bener, Taylor Simao, Abigail E. Descoteaux, Boris M. Slepchenko, Mayu Inaba

## Abstract

Stem cell niche signals act over a short range so that only stem cells but not the differentiating daughter cells receive the self-renewal signals. *Drosophila* female germline stem cells (GSCs) are maintained by short range BMP signaling; BMP ligands Dpp/Gbb activate receptor Tkv to phosphorylate Mad (phosphor-Mad or pMad) which accumulates in the GSC nucleus and activates the stem cell transcription program. pMad is highly concentrated in the nucleus of the GSC, but is immediately downregulated in the nucleus of the pre-cystoblast (preCB), a differentiating daughter cell, that is displaced away from the niche. Here we show that this asymmetry in the intensity of pMad is formed even before the completion of cytokinesis. A delay in establishing the pMad asymmetry leads to germline tumors through conversion of differentiating cells into a stem cell-like state. We show that a Mad phosphatase Dullard (Dd) interacts with Mad at the nuclear pore, where it may dephosphorylate Mad. A mathematical model explains how an asymmetry can be established in a common cytoplasm. It also demonstrates that the ratio of pMad concentrations in GSC/preCB is highly sensitive to Mad dephosphorylation rate. Our study reveals a previously unappreciated mechanism for breaking symmetry between daughter cells during asymmetric stem cell division.

## Introduction

Stem cells often divide asymmetrically to generate a stem cell and a differentiating daughter cell(Morrison and Kimble, 2006). It remains poorly understood how a stem cell and a differentiating daughter cell can receive distinct levels of signal and thus acquire different cell fates (self-renewal vs. differentiation) despite being adjacent to each other and thus seemingly exposed to similar levels of niche signaling.

The *Drosophila* female germline stem cell (GSC) is an excellent model to study niche-stem cell interaction because of its well-defined anatomy and abundant cellular markers(Spradling et al., 2011). At the tip of each ovariole, two to three GSCs adhere to the cluster of niche cells, known as the cap cells (CCs) (Figure 1A). A bone morphogenetic protein (BMP) ligand, Decapentaplegic (Dpp), is secreted by the CCs and is the essential factor for GSC maintenance (Xie and Spradling, 1998) (Song et al., 2004). Dpp binds to the serine-threonine kinase receptor Thickveins (Tkv) expressed on GSCs. Activated Tkv then phosphorylates Mothers against decapentaplegic (Mad) to create phosphorylated Mad (pMad) that enters the nucleus and directly binds to the promotor of the differentiation factor *bam* to downregulate transcription. The downregulation of *bam* is essential for GSC self-renewal (Xie and Spradling, 1998) (Song, 2004; Akbar et al., 2009; Chen and McKearin, 2003a). Multiple studies have defined mechanisms for fine-tuning Dpp signal strength and range (Xia et al., 2012; Wang et al., 2008; Van De Bor et al., 2015; Wilcockson and Ashe, 2019; Guo and Wang, 2009; Harris et al., 2011; Liu et al., 2010; Schulz et al., 2002; Xia et al., 2010; Jiang et al., 2008; Tseng et al., 2018). A well-documented example is the discrimination of GSC and preCB, a differentiating daughter cell, by Fused (Fu), a serine/threonine kinase. Fu phosphorylates Tkv and promotes its ubiquitination by E3 ligase Smurf and thus subsequent degradation (Xia et al., 2010). On the other hand, pMad promotes the degradation of Fu; therefore, Fu is concentrated only in the preCB. These mechanisms initiate a feedback loop to enhance the gradient of BMP response downstream of the niche signal (Xia et al., 2012). However, to date, how Mad, the immediate downstream molecule of the niche BMP ligand, is initially regulated during GSC division has not been studied.

**Figure 1.**
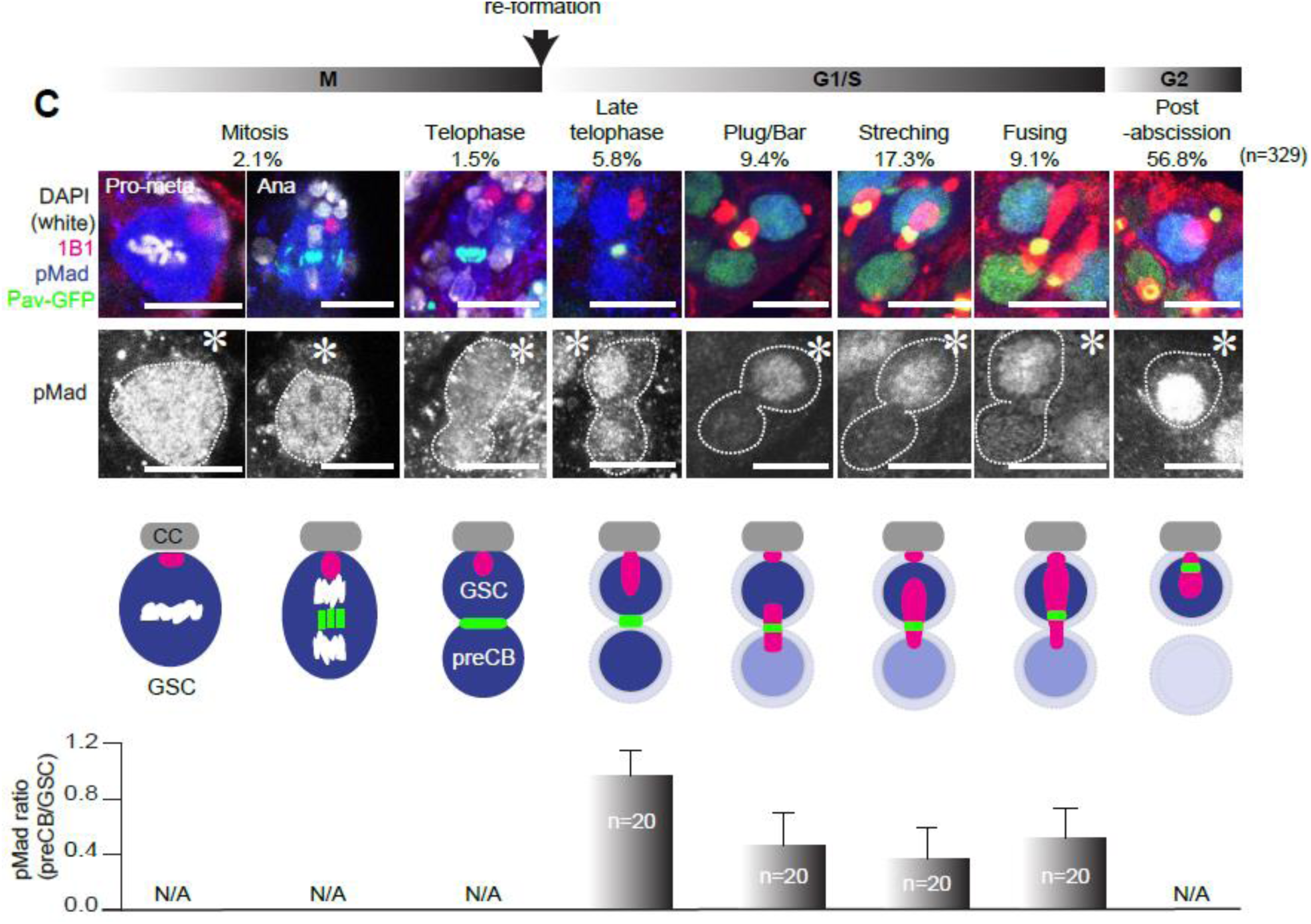
GSC and preCB establish asymmetric pMad levels prior to the completion of abscission. A) Schematic of the *Drosophila* female GSC niche. B) A representative confocal image of the GSC niche with an GSC/preCB pair. 1B1 (red, fusome), Pav-GFP (green, contractile ring) and pMad (blue). C) Measured pMad ratios (preCB/GSC) during cell cycle stages. Cell cycle stages were assessed based on previously documented fusome morphology and the location of the contractile ring (Pav-GFP). G1/S phase was subdivided into the following phases; Plug/Bar, Stretching, Fusing. Percentages of each stage are shown in round brackets (329 GSCs were scored). Representative confocal images of each stage are shown in the upper panel. The pMad channel is shown below (Grayscale). Schematics of GSC/preCB pair of each stage 1B1 (pink, fusome), Pav-GFP (green, contractile ring) and pMad (blue). The graph at the bottom shows the ratios of nuclear pMad intensities (preCB’s divided by GSC’s). pMad intensities were measured and background from each sample was subtracted from each measurement. Asterisks indicate the location of CCs. Scale bars, 10 μm.

In this study, we demonstrate that pMad rapidly reaches different levels in dividing GSCs after the mitosis but before the completion of cytokinesis. It remains high in the nuclei of future GSCs, while decreasing in the nuclei of the preCBs. After mitosis, the GSC takes several hours to complete cytokinesis (Xia et al., 2012; de Cuevas and Spradling, 1998; Gilboa et al., 2003; Chen and McKearin, 2003b). During this time, the preCB still shares its cytoplasm with the GSC (Matias et al., 2015). It is unclear how the GSC and the preCB can have different levels of pMad in their nuclei. We show that the previously known Mad phosphatase Dullard (Dd) plays an essential role in formation of a sharp boundary between GSC and preCB (i.e. high vs. low pMad). Although Dd itself does not exhibit asymmetric localization in GSC and preCB, mathematical modeling shows that biased phosphorylation of pMad near the niche (due to local activation of Tkv on the niche side) combined with unbiased dephosphorylation by equally distributed Dd is sufficient to explain the observed pMad asymmetry. In summary, our results provide a mechanism by which a self-renewal program is confined to the stem cells during asymmetric division.

## Results

### GSC and preCB establish asymmetric pMad levels prior to the completion of abscission

After mitosis, GSC/preCB pairs maintain their cytoplasmic connection almost throughout G1/S phase(Matias et al., 2015). When mEOS-αTub expressed in germ cells (nos> mEOS-αtub) was photoconverted in either GSC only or preCB only, the converted signal of cytoplasmic mEOS-αTub quickly equalized between GSC and preCB (120s ± 21.7, n=18, Figure S1A). Equilibration of the signal was observed for both directions, GSC to preCB or preCB to GSC (Figure S1A). Consistent with a previous study(Matias et al., 2015), the cytoplasmic connectivity judged by rapid exchange of photoconverted mEOS-αTub lasted until the completion of GSC-preCB abscission (40% of GSCs; 24 GSCs were judged as “connected” out of n=60 GSCs tested). Similarly, GFP tagged Mad protein (GFP-Mad) also showed equalization after photobleaching (Mad: 99.3s ± 16.7, n=15) (Figure S1B, C, video1), indicating that Mad can freely diffuse between GSC and preCB, and no detectable immobilized fraction of Mad was observed. These data also suggest that Mad likely travels from the plasma membrane to the nucleus largely by a diffusion-based mechanism.

Despite the rapid exchange of Mad in GSC/preCB pairs in G1/S phase, we observed that pMad signaling became clearly asymmetric between GSC and preCB nuclei soon after telophase, but well before the completion of cytokinesis (Figure 1A and 1B). In a mitotic GSC, pMad distributed uniformly within the cell (Figure 1C, left 3 panels). In telophase cells, pMad exhibited a similar level of nuclear signal intensity between GSC and preCB (Figure 1C, Late telophase). However, soon after telophase, the pMad level became clearly asymmetric between GSC and preCB, much earlier than the completion of cytokinesis (Figure 1C, Plug/Bar). We therefore conclude that GSC/preCB pairs establish pMad asymmetry while they freely exchange Mad protein (Figure S1A), suggesting that asymmetric accumulation of pMad between GSC and preCB nuclei is not due to a diffusion barrier between these two cells. These data prompted us to investigate the cellular mechanism by which pMad is concentrated more in GSC nuclei than preCB nuclei, breaking the symmetry of niche signaling between GSCs and preCBs.

### Dd is required for establishment of pMad asymmetry through dephosphorylating Mad at the nuclear pore

We performed a candidate RNAi mini screening to identify the genes that affect pMad asymmetry. We identified a phosphatase, Dullard (Dd) as being required for pMad asymmetry between GSC and preCB. Dd is a phosphatase that has been shown to suppress BMP signaling by directly dephosphorylating Mad in *Drosophila* wings and *Drosophila* S2 cell lines (Liu et al., 2011; Urrutia et al., 2016). To test the function of Dd in pMad asymmetry formation, we knocked down Dd in the germline (nos>dd RNAi; the knock down efficiency was validated as described in the *Methods*). Dd knocked down GSC showed elevated pMad intensity (normalized by Mad-GFP intensity), suggesting that Dd is the major Mad phosphatase in female GSCs (Figure 2A-C). Moreover, without Dd, the ratio of the pMad intensity in the GSC nucleus to that in the preCB nucleus was close to 1 (symmetric) until cytokinesis was complete (Figure 2D, E), indicating that Dd is necessary for early establishment of pMad asymmetry. Although Dd was originally identified as a nuclear envelope phosphatase required for nuclear envelope integrity via dephosphorylating other substrates than Mad (Kim et al., 2007), we did not observe aberrant nuclear envelope morphology in Dd mutant cells as shown in other systems (Kim et al., 2007) (Figure S2), suggesting that Dd likely regulates pMad asymmetry via directly dephosphorylating Mad.

**Figure 2.**
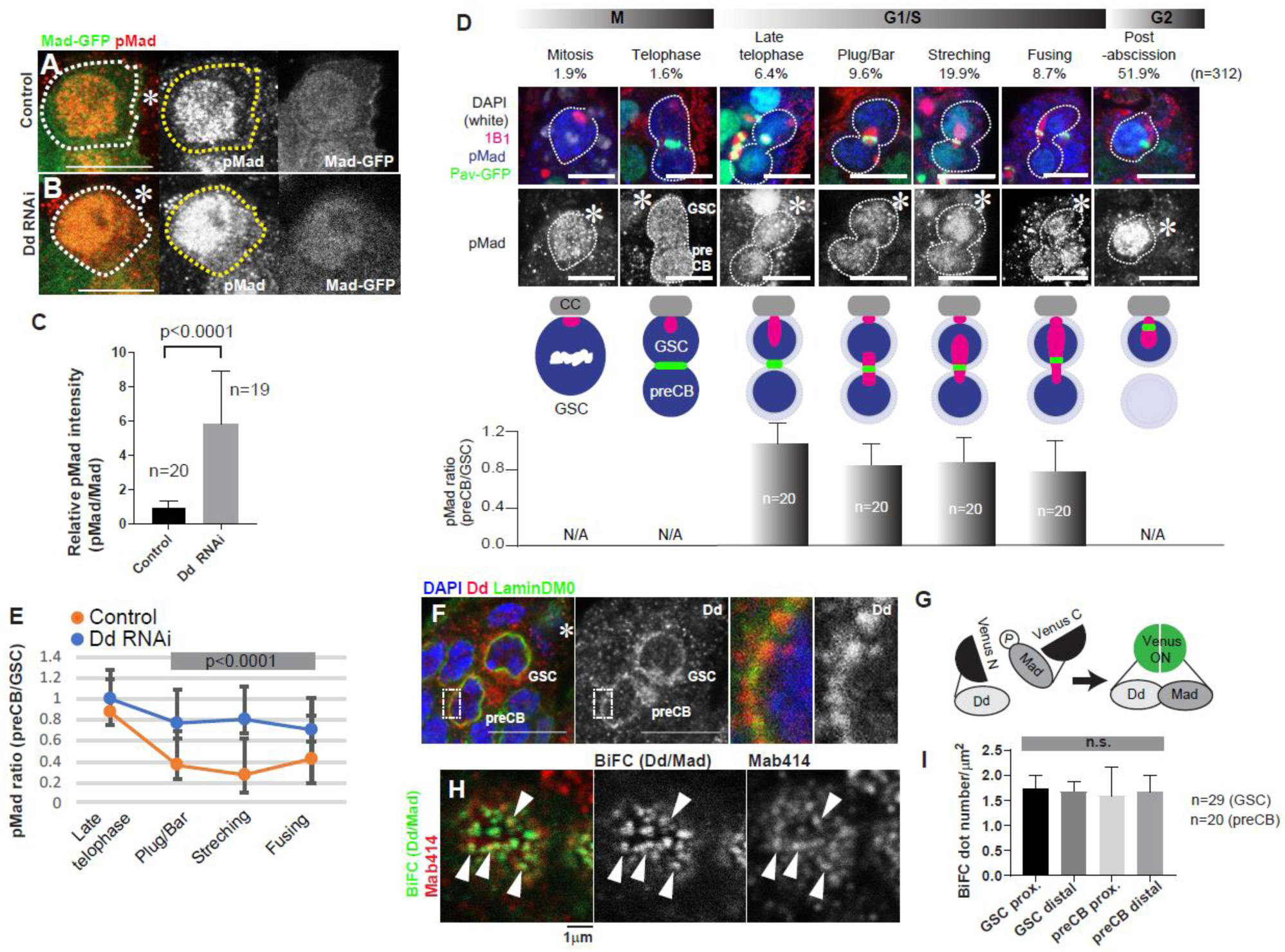
Dd is required for pMad asymmetry formation. A-C) Comparison of relative pMad intensities (pMad/GFP-Mad) in GSCs with or without Dd RNAi. D) Representative images of pMad staining during cell cycle stages of GSCs in Dd RNAi ovary (nos>dd RNAi). Graph in the bottom shows the ratio of nuclear pMad intensities (preCB’s divided by GSC’s, see Figure1C legend). NA: not applicable. E) Comparison of pMad ratio (preCB/GSC) during GSC division with or without Dd RNAi. F) A representative image of co-staining of Dd (nos>dd-VNm9, anti GFP staining, Red) and nuclear envelope marker, Lamin DM0 (Green). DAPI; Blue. Regions within square are magnified in the right two panels. G) Schematic of Mad-Dd BiFC design. N-terminus half and C-terminus half of Venus were fused to the N-terminus of Dd cDNA or the C-terminus of Mad cDNA, respectively. When these constructs are expressed together, Venus is reconstituted upon Dd-Mad interaction and emits fluorescence. H) The Dd-Mad BiFC signal was observed as spots on the nuclear membrane. Arrowheads show co-localization of the BiFC signal and the pan-nuclear pore marker, Mab414 staining. I) Nuclear pore number/area scored by Mab414 staining did not show bias throughout of GSC and preCB nuclei. Prox (proximal half nuclear envelope), Distal (distal half nuclear envelope). Approximately 2×2μm of nuclear surface regions (from the indicated number of GSCs) were scored for each data point. Asterisks indicate the location of CCs. Scale bar, 10 μm. ns: non-significant (P≥0.05).

Dd has been shown to localize to the nuclear envelope. Consistently, we found that Dd protein expressed in GSCs localizes along the Lamin DM0, a nuclear envelope marker, in GSCs (Figure 2F: determined by an anti-GFP antibody that recognizes the N terminal half of Venus (VNm9) (Saka et al., 2007) tagged Dd (Urrutia et al., 2016; Kim et al., 2007). A bimolecular fluorescence complementation (BiFC) assay showed that Mad and Dd interact specifically at the nuclear pore. When the C terminus of Dd was fused to VNm9 and the C terminus of Mad was fused with the C-terminal half (VC) of Venus fluorescent protein (Figure 2G) (Saka et al., 2007), a strong BiFC signal was observed as a punctate pattern along the nuclear envelope (Figure 2H and I). The BiFC signal was co-localized with nuclear pore marker Mab314 (Figure 2J). These data indicate that Dd and Mad interact at the nuclear pore, and imply that Dd may directly phosphorylate pMad when Mad shuttles between the nucleus and cytoplasm. Together, our results show that Dd, a phosphatase known to dephosphorylate Mad, plays a critical role in forming asymmetry in pMad intensity between GSC and preCB.

How does Dd contribute to establishing pMad asymmetry? Dd distribution was not asymmetric between GSC and preCB (Figure 2F), suggesting that asymmetric localization/level of Dd is not the cause of the asymmetric pMad level. Likewise, we observed no biased distribution of Mab414/BiFC positive nuclear pores between GSC and preCB (Figure 2I).

### Modeling suggests that pMad asymmetry emerges due to the interplay of phosphatase activity and pMad diffusion

The Dpp ligand is reported to be highly concentrated and stabilized on the surface of CCs; Wilcockson and Ashe, 2019; Guo and Wang, 2009; Liu et al., 2010, Entchev et al., 2000; Akiyama et al., 2008). A recent study demonstrated that GSCs project cellular protrusions into CCs to access a reservoir of Dpp (Wilcockson and Ashe, 2019). On the other hand, somatic escort cells (ECs) surrounding GSCs express Tkv receptor to absorb any free Dpp(Luo et al., 2015). These mechanisms possibly constrain Tkv activation to the side of GSC near the niche.

Our mathematical modeling showed that asymmetric Tkv/Dpp and equal distribution of Dd can explain pMad asymmetry. The model includes essential processes affecting the spatiotemporal distribution of pMad in a dividing stem cell (Figure 3A): (i) phosphorylation of Mad by activated kinase receptors residing on the plasma membrane at the pole of GSC; (ii) diffusion of Mad and pMad throughout the shared cytoplasm, (iii) shuttling of pMad and Mad between the cytosol and nuclei via nuclear pores; (iv) binding of pMad in the nuclei of GSC and preCB to the nuclear matrix and DNA; (v) dephosphorylation of free pMad by phosphatases in the inner leaflet of the nuclear envelope; and (vi) diffusion of free pMad and Mad in the nuclei. Except for the phosphorylation event occurring only at one pole of GSC, identical parameters were used to characterize the same processes in both GSC and preCB. The model was solved in a three-dimensional (3D) geometry mimicking a shape of a cell undergoing cytokinesis (Figure 3B; see subsection ‘Model geometry’ of ‘Mathematical modeling’ in *Methods*). The kinase (activated Tkv receptor) is assumed to be concentrated within a ‘cap’ shown in red in Figure 3C. Model parameters are listed in Figure 3D (see ‘Model parameters’ under ‘Mathematical modeling’ in *Methods*).

**Figure 3.**
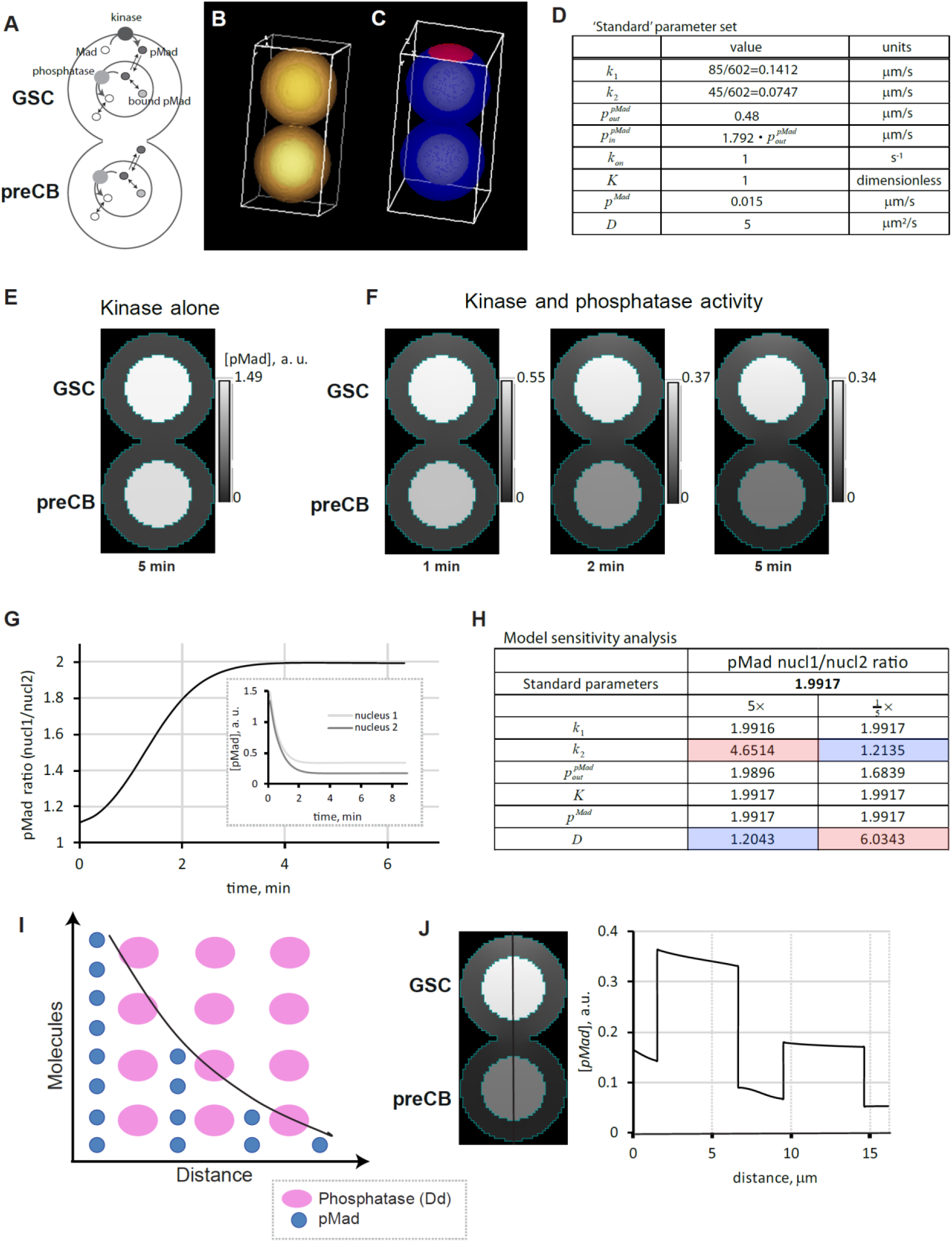
pMad asymmetry is determined by the interplay of phosphatase activity and pMad diffusion. A) Fluxes and reactions essential for partitioning of pMad between daughter cells. B) Model geometry. C) Simulated localization of the kinase (shown in red). D) ‘Standard ‘parameter set constrained by experimental data. The parameters are defined as follows: *k*_1_ is the kinase activity constant; *k*_2_ is the phosphatase activity constant, 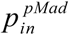 is the nuclear envelope permeability to pMad import; 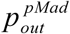 is the nuclear envelope permeability to pMad export; *k*_*on*_ is the rate constant of pMad binding to the nuclear matrix and DNA, and *K* is the corresponding dissociation equilibrium constant; *p*^*Mad*^ is the nuclear envelope permeability to pMad import/export; *D* is the pMad diffusivity (see also *Methods*). E) Distribution of pMad after phosphorylation of Mad for 5 minutes in the absence of phosphatase activity. F) Onset of asymmetry of pMad partitioning after the phosphatase ‘is turned on’ and the phosphorylation and dephosphorylation occur simultaneously. G) Time course of the ratio of the pMad concentrations in the nuclei; time zero corresponds to the time when dephosphorylation begins. Inset: time courses of pMad concentrations in the nuclei of GSC and preCB. H) Model sensitivity analysis indicates strong correlation between the degree of pMad asymmetry and the rate of phosphatase activity (for parameter definitions, see the legend for panel D and *Methods*; for values, see panel D). I) Diagram of a simplified 1D model, where pMad (blue circles), produced at a left boundary, diffuses throughout and is dephosphorylated by a phosphatase (pink ovals) evenly distributed throughout the ‘cell’. J) Vertical line scan of pMad steady-state concentrations in the 3D model. Left panel: pMad distribution in the 3D model with ‘standard’ parameters (D) at *t* = 1000 s.Right panel: the pMad concentration as function of the distance counted from the top along the vertical line shown in the left panel.

Given that the active kinase (Dpp bound Tkv) localizes only at one pole of the GSC, asymmetric partitioning of pMad between the GSC and preCB may appear intuitive. However, our model yields nearly equal steady-state concentrations of pMad in the nuclei of GSC and preCB, if the phosphatase activity is relatively low or zero. It is only when the phosphatase activity is sufficiently high, that the substantial asymmetry of the pMad partitioning becomes apparent. Figure 3E-G illustrates results of a numerical experiment where Mad, initially unphosphorylated and uniformly distributed throughout the cells in the cytoplasm and nuclei is first subjected to kinase activity alone for five minutes, after which the phosphatase ‘turns on’ and both phosphorylation and dephosphorylation occur simultaneously.

Phosphorylating Mad in the absence of phosphatase activity yields a ratio of pMad concentrations in the nuclei of GSC and preCB close to unity (Figure 3E, 3G). The asymmetry of pMad in the daughter stem cells emerges in the model as a result of phosphatase activity, as demonstrated in Figure 3F, with the ratio of the nuclear concentrations of pMad approaching 2:1 (Figure 3G). The inset in Figure 3G depicts the corresponding time courses of pMad concentrations in the two nuclei. Thus, the model reproduces our experimental results.

We next analyzed the model to determine the factors that have the most impact on the pMad nucl1/nucl2 ratio (here and below ‘nucl1’ and ‘nucl2’ stand for nucleus 1 and nucleus 2 denoting the nuclei of GSC and preCB, respectively). Figure 3H shows sensitivities of this ratio at steady state to a 5-fold increase/decrease of ‘standard’ parameters listed in Figure 3D. The results in Figure 3H show that pMad diffusivity is the only other parameter whose effect on the pMad asymmetry is comparable to that of the phosphatase activity constant (for parameter notation and definitions, see ‘Mathematical modeling’ in Methods). Therefore, the pMad nucl1/nucl2 ratio is largely determined by the interplay of phosphatase activity and pMad diffusion.

The results in Figure 3E-H were obtained on the assumption that no diffusion barrier exists between GSC and preCB before the completion of cytokinesis. Because the existence of such a barrier in the connected GSC/preCB pair would affect the partitioning of pMad, we examined a model where the barrier between the GSC and preCB is formed before the cytokinesis is completed. The steady state of this model (Figure S3A) is notably different from the pMad partitioning observed in the majority of visibly connected GSC/preCB pairs. Rather, it is characteristic of what occurs after the completion of cytokinesis, when the pMad intensity in GSCs becomes significantly higher than in the connected GSC/ preCB pairs (Figure 1C, G2 phase GSC, Figure S3B) and the pMad signal in preCB essentially disappears. Indeed, consistent with these observations, the simulated pMad concentration quickly increases in the GSC and decreases in the preCB after the barrier is formed (Figure S3A, upper panel). These results are thus in agreement with the experimental data described in section ‘GSC and preCB establish asymmetric pMad levels prior to the completion of abscission’ that indicate the absence of a diffusion barrier in the majority of the visibly interconnected GSC/preCB pairs.

### The underlying mechanism: origins of pMad gradients and their steepness

The simulation results described in the previous section can be elucidated further by noting that in our system, kinases and phosphatases localize differently: Tkv receptors reside on the plasma membrane, whereas Dd molecules co-localize with nuclear pores. The separation of pMad ‘sources’ and ‘sinks’ in space produces steady-state pMad gradients, whose steepness depends on the dephosphorylation of pMad, which makes them steeper, and on the pMad diffusion that levels them out.

This can be illustrated by a simple one-dimensional (1D) model, where pMad, produced by a kinase at a ‘cell’ boundary, diffuses into the ‘cell’ interior, where it is dephosphorylated by a phosphatase evenly distributed throughout the ‘cell’ (Figure 3I). In a long 1D ‘cell’, a distribution of the pMad concentration, [*pMad*](*x*), is well approximated by a descending exponential (the black curve in Figure 3I), described by [*pMad*](*x*) =[*pMad*](0)exp(−*x*/ *λ*), where *x* denotes a location in the 1D cell and *λ* is the length parameter, determined exclusively by the pMad diffusivity (*D*) and phosphatase activity constant 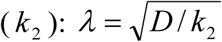 (Because the phosphatase in this model is distributed throughout the cell, the units of *k*_2_ are s^-1^). Note that the pre-exponential factor is determined by kinase activity (*k*_1_).

Parameter *λ* characterizes the steepness of pMad gradient. Indeed, the pMad ratio computed in the 1D model for two points separated by a given distance *d* is exp(*d* / *λ*), so it is fully determined by *λ*, i.e. only by *k*_2_ and *D*. In the realistic 3D model, the pMad cytosolic concentrations are ‘sampled’ by the nuclei, where they are also amplified due to pMad binding inside the nuclei and unequal permeabilities of the nuclear envelope to inward/outward pMad fluxes. Figure 3J presents a line scan along the vertical axis of the steady-state concentrations of pMad obtained with the ‘standard’ parameters (Figure 3D).

The simplified 1D model also explains why the pMad ratio does not depend on kinase activity. Varying the phosphorylation rate does change pMad concentrations, but they all change by the same factor that cancels out in the ratio.

Note that according to the 1D model, the pMad ratio becomes large if *λ* < *d*, or 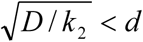,or 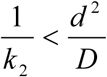. In other words, a large pMad ratio is achieved if the time required to dephosphorylate a pMad molecule is shorter than the time it takes the pMad molecule to diffuse a distance separating the two nuclei. Because in the 3D model, the dephosphorylation occurs on the inner leaf of the envelope, the effective dephosphorylation time also includes the time of crossing the envelope, which depends on the envelope permeabilities. This explains some sensitivity of the pMad ratio to *p*^*pMad*^ (Figure 3H).

In summary, the model reproduces the experimentally observed asymmetric accumulation of pMad during *Drosophila* female GSC division. The model also shows that this is a robust phenomenon originating from spatial separation of kinases and phosphatases: the kinases reside on the plasma membrane, whereas the phosphatases localize to the nuclear envelopes. This brings about steady-state gradients of pMad in the cytoplasm, whose steepness, and therefore the nucl1/nucl2 ratio, is a function of the effective dephosphorylation rate and pMad diffusivity and is independent of other model parameters.

### pMad asymmetry is required for GSC to preCB fate transition

To understand the biological significance of Dd-mediated establishment of pMad asymmetry, we examined the Dd mutant phenotype. A Dd hypomorphic mutant exhibited a slightly increased number of pMad positive, GSC-like cells near the niche (3.3±1.5 (n=27) in control, 4.6±1.2 (n=27) in dd mutant, p<0.001, Figure 4A, B). Expression of constitutively active Tkv receptor (TkvCA, active kinase activity without ligand binding) is often utilized to induce the overproliferation of undifferentiated germ cells outside of the niche (referred to as germ cell tumors hereafter) (Casanueva and Ferguson, 2004). We found that these tumors showed only weak pMad staining (Figure 4C), likely because the TkvCA is subjected to protein degradation as is known for endogenous Tkv protein (Xia et al., 2012; Xia et al., 2010). Consistent with the previous studies, these tumor cells possessed round fusomes, the hallmark of undifferentiated cells. Combined with weak pMad staining, these tumor cells resemble preCB (round fusome, low pMad) (Figure 4C, E) (Jiang et al., 2008) (Casanueva and Ferguson, 2004). However, introducing the Dd hypomorphic allele (*ddd*^*P*^) to the TkvCA background led to strong pMad in tumor cells (Figure 4D). High pMad and a round spectrosome indicates GSC identity, and this condition (*ddd*^*P*^ /+ and TkvCA) significantly increased proliferation of tumor cells (Figure 4F, G), possibly due to cell fate switching from a preCB to a GSC-like state (Figure 4H). Taken together, our study shows that downregulation of niche signaling before stem cell and its differentiating daughter cell complete abscission plays an important role to promote differentiation and prevent tumor formation as differentiating daughters exit the stem cell niche.

**Figure 4.**
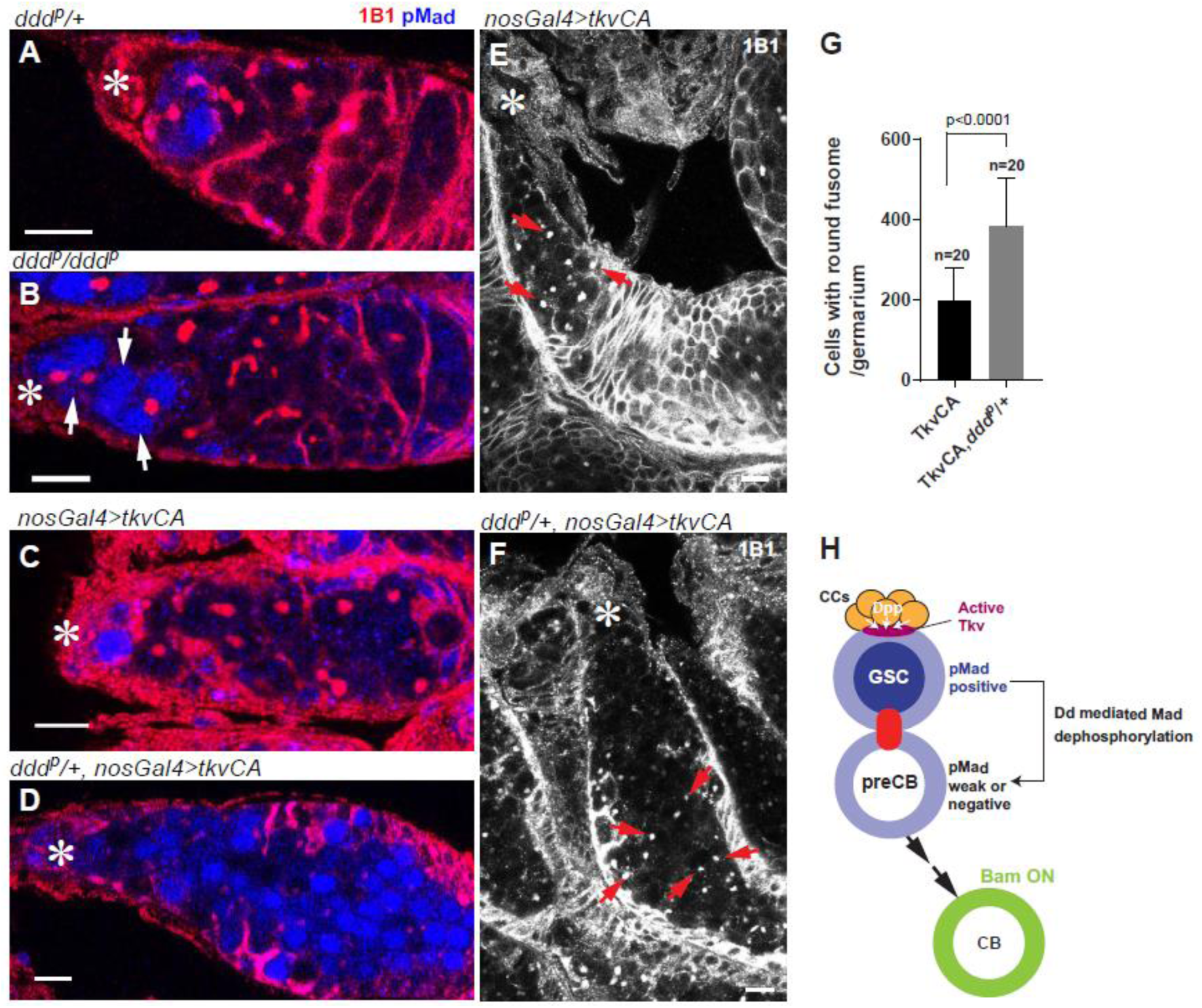
Introducing a heterozygous Dd mutation increased pMad positive cells within TkvCA tumor and increased the tumor size. A-D) 1B1(spectrosome, fusome, red) and pMad staining (blue) of germarium from indicated genotypes. B) White arrows indicate extra pMad positive cells away from CCs. C, D) pMad staining of TkvCA expressing tumor cells without (C) or with (D) introducing one copy of *ddd*^*p*^ mutant allele. E, F) Examples of typical germarium images of TkvCA expressing tumor without (E) or with (F) one copy of *ddd*^*p*^ mutant allele. E) Red arrows show round fusomes, a hallmark of undifferentiated cells. G) Comparison of tumor cell number (cells with round fusome) in TkvCA expressing germarium with or without *ddd*^*p*^ mutant allele. 20 germaria were scored for each data point. H) Model. pMad asymmetry formation ensures GSC to preCB differentiation. Asterisk indicates the location of CCs. Scale bars: 10 μm.

## Discussion

During asymmetric stem cell division, two daughter cells acquire different cell fates. The *Drosophila* ovarian niche differentially activates GSC and preCB due to the distinct position of these cells, one adjacent to the niche, the other displaced from the niche. What is the initial cause of the difference? While many factors have been identified that can amplify and/or ensure the already existing niche signal difference, it has been unknown how Mad, the immediate downstream molecule of the niche BMP ligand, is initially regulated during GSC division.

Upon the activation of the niche signal receptor Tkv kinase, its substrate, Mad, is phosphorylated near the plasma membrane and then travels throughout the cytoplasm toward the nucleus. Our study showed that this occurs largely via a diffusion-based mechanism. GSC-preCB pairs continue to share cytoplasm for at least several hours after mitosis, throughout almost the entire G1-S phase (Matias et al., 2015). Consistently with previous studies, we observed that Mad protein diffuses throughout the cytoplasm of the GSC and preCB. We found however, that during this phase, the pMad levels in the nuclei of the GSC and preCB are already asymmetric. Our candidate approach determined that the local activation of the kinase at the niche-GSC contact site is not sufficient for asymmetric pMad partitioning, and discovered that the Mad phosphatase, Dd, that dephosphorylates Mad at the nuclear pores in GSC and preCB alike, is the essential factor for the formation of early asymmetric partitioning of pMad.

To gain insight into mechanisms responsible for the asymmetric distribution of pMad between daughter cells, we formulated a mathematical model, which included all major factors governing the spatiotemporal dynamics of pMad, and constrained it by the experimental data obtained in this study. Our model suggests that the localization of the activated kinase to the site of contact of the GSC plasma membrane and the niche and the symmetric distribution of the phosphatase between the nuclear envelopes of the GSC and preCB are sufficient to explain the experimentally observed asymmetry of the pMad levels in the GSC and preCB.

Analysis of the model revealed that for the pMad asymmetry to occur, it is necessary that the positive and negative regulators of pMad (the kinases and phosphatases) be separated in space, as this brings about spatial gradients of pMad. However, this condition is not sufficient, as the gradients could be shallow. The modeling showed, furthermore, the steepness of pMad gradients is exclusively determined by the interplay of two factors: pMad dephosphorylation that steepens the gradient, and pMad diffusion in the cytoplasm that levels it out. Active regulation of the pMad diffusivity, which is determined by protein size and the effective viscosity of the cytoplasm (Berg, 1993), is unlikely. This makes the rate of pMad dephosphorylation the single most important parameter determining the pMad asymmetry. In turn, the phosphatase activity is the essential factor that determines this rate. The effective dephosphorylation rate also depends, to some degree, on the permeability of nuclear pores to pMad import, because dephosphorylation of pMad occurs inside the nucleus.

Our study identifies and explains a mechanism by which a stem cell can rapidly set up the initial asymmetry with respect to an extrinsic signal, providing a conceptual framework for understanding the dynamics of niche-stem cell signaling.

## Acknowledgements

We thank Yasushi Saka, Edward Eivers, Shin Sugiyama, the Bloomington *Drosophila* Stock Center and the Developmental Studies Hybridoma Bank for reagents; Michael Buszczak, Yukiko Yamashita, Mark Terasaki and Laurinda Jaffe for discussion. Christopher Bonin and Luisa Lestz for manuscript editing. B.M.S. thanks Leslie Loew for continuing support. This research is supported by 1R35GM128678-01 from National Institute for General Medical Sciences and startup funds from UConn Health (to M.I.) and by P41GM103313 from National Institute for General Medical Sciences. The Virtual Cell is currently supported by R24 GM13421 from the National Institute for General Medical Sciences.

## Author Contributions

M.I. conceived the project, designed and executed experiments and analyzed data. J.S., M.B.B., T.S. and A.D. executed experiments and analyzed data. B.M.S. formulated the mathematical model, ran simulations and analyzed simulation results. All authors wrote and edited the manuscript.

## Declaration of Interests

The authors declare no competing interests.

## Methods

### Fly husbandry and strains

All fly stocks were raised in standard Bloomington medium at 25°C. The following fly stocks were used, UASp-tkvCA(Guo and Wang, 2009) (gift from Michael Buszczak), Ubi-Pavarotti (Pav)-GFP(Minestrini et al., 2002) (gift from Yukiko Yamashita), hypomorphic Dd mutants (*ddd*^*P*^) (gift from Shin Sugiyama)(Liu et al., 2011). The following stocks were obtained from the Bloomington stock center: UAS-mEOS-αtub (BDSC51314); for Dd RNAi, short hairpin RNA (TRiP.GL01268, BDSC41840) was expressed under the control of nosGal4 (see below for validation method). Control cross for RNAi was designed with matching gal4 and UAS copy number using TRiP background stocks (Bloomington Stock Center BDSC36304 or BDSC35787) at 25 °C.

### Generation of pUASp-dd-VNm9, pUASp-VC-mad and pUASp-GFP-mad flies

#### pUASp-dd-VNm9

N-terminal half of VENUS with point mutations (VNm9) (Saka et al., 2007) fragment was amplified from the synthesized gBlock fragment (VNm9 gBlock) using VNm9-F and VNm9-R primers.

The Not1-Kozak-dd-BglII fragment was amplified by using NotI-Kozak-dd-F, BglII-dd-R primers. The pUASp-attB vector was digested with NotI and AscI, then ligated with Not1-Kozak-dd-BglII and BglII-Linker-VNm9-Asc1 fragments.

Transgenic flies were generated using strain attP2 by PhiC31 integrase-mediated transgenesis (BestGene).

#### pUASp-GFP-mad

BglII NotI Mad-F and AscI STOP Mad-R primers were used to amplify wild type Mad cDNA (RA) from a testis cDNA pool. Then, the NotI-GFP-NotI fragment was amplified from the pPGW vector (https://emb.carnegiescience.edu/drosophila-gateway-vector-collection#_Copyright,_Carnegie) using NotI GFP-F and NotI GFP-R primers and inserted into the NotI site located in the BglII NotI Mad-F primer. Transgenic flies were generated using strain attP40 by PhiC31 integrase-mediated transgenesis (BestGene).

#### pUASp-VC-mad

BglII NotI Mad-F and AscI STOP Mad-R primers were used to amplify wild type mad cDNA (RA) from a testis cDNA pool, then inserted into pUASp-attB vector. Not I VC155-F and NotI linker VC155-R primers were used to amplify the C-terminal (VC) half of VENUS from pCS2+ VC (Saka et al., 2007) (gift from Y. Saka). The NotI-VC155-NotI fragment was inserted into the NotI site located in the mad forward primer. Transgenic flies were generated using strain attP40 by PhiC31 integrase-mediated transgenesis (BestGene).

### Immunofluorescent staining

Ovaries were dissected in phosphate-buffered saline (PBS) and fixed in 4% formaldehyde in PBS for 30–60 minutes. Next, ovaries were washed in PBST (PBS +0.1% Tween 20) for at least 30 minutes, followed by incubation with primary antibody in 3% bovine serum albumin (BSA) in PBST at 4 °C overnight. Samples were washed for 60 minutes (three times for 20 minutes each) in PBST, incubated with secondary antibody in 3% BSA in PBST at 4 °C overnight, and then washed for 60 minutes (three times for 20 minutes each) in PBST. Samples were then mounted using VECTASHIELD with 4′,6-diamidino-2-phenylindole (DAPI). The primary antibodies used were in Key Resources Table. AlexaFluor-conjugated secondary antibodies were used at a dilution of 1:400. Images were taken using a Zeiss LSM800 confocal microscope with a 63 ×oil immersion objective (NA=1.4) and processed using Fiji. Further details are available in (Inaba and Yamashita, 2017).

### Live imaging

Ovaries from newly eclosed flies were dissected in 1 ml of prewarmed Schneider’s *Drosophila* medium supplemented with 10% fetal bovine serum and glutamine–penicillin–streptomycin. Dissected ovaries were placed onto ‘Gold Seal™ Rite-On™ Micro Slides two etched rings’ with a drop of media, then covered with coverslips. An inverted Zeiss LSM800 confocal microscope with a 63 ×oil immersion objective (NA=1.4) was used for imaging.

### Photo-conversion and photo-bleaching

Photo-conversion of mEOS-αTub or photo-bleaching of GFP-Mad was accomplished using a Zeiss LSM800 confocal laser scanning microscope with 63X/1.4 NA oil objective. Zen software was used for programming of each experiment. Laser powers and iteration were optimized to achieve an approximately 50%-70% conversion or an approximately 70%-100% bleach; a 405 nm laser (photoconversion) and a 480nm laser (photocbleaching) were used. Fluorescence recovery was monitored every 10 seconds for up to 10 minutes.

### Quantification of pMad ratio

Ovaries from RNAi flies expressing Pav-GFP (midbody ring) were dissected and stained with 1B1 (fusome) and Vasa (germ cell cytoplasm) to identify interconnected GSC/CB pairs. To calculate the ratio of nuclear pMad between GSC and preCB, integrated intensity within the GSC nuclear region was measured for anti-pMad staining and divided by the area and then background obtained from the same field was subtracted.

### Quantitative reverse transcription PCR to validate RNAi-mediated knockdown of genes

Females carrying nosGal4 driver were crossed with males of the dd RNAi line. Ovaries from 20 female progeny, age 3-7 days, were collected and homogenized by pipetting in TRIzol Reagent (Invitrogen) and RNA was extracted following the manufacturer’s instructions. One microgram of total RNA was reverse transcribed to cDNA using SuperScript III First-Strand Synthesis Super Mix (Invitrogen) with Oligo (dT)20 Primer. Quantitative PCR was performed, in duplicate, using SYBR green Applied Biosystems Gene Expression Master Mix on an CFX96 Real-Time PCR Detection System (Bio-Rad). Relative quantification was performed using the comparative CT method (ABI manual). ddRNAi reduced dd transcript to 23.9% of control.

### Mathematical modeling

#### Model geometry

Cells are defined by two intersecting identical spheres with an 8.3-μm diameter. Two nuclei are modeled by spheres with 5-μm diameters that are concentric with respective large spheres.

Parameters of the model geometry are the averages of the respective sizes of measurements from experimental images (n=15 for GSCs and n=15 for preCB).

Let *Ω*_*cyto*_, *Ω*_*nucl*1_, and *Ω*_*nucl*2_ denote the spaces occupied by the cytosol and nuclei of GSC and CB, respectively, and (*x, y, z*) are the Cartesian coordinates of a spatial point. The nuclei *Ω*_*nucl*1_ and *Ω*_*nucl*2_ are then modeled as {*Ω*_*nucl*2_ | (*x* ^2^ + (*y* − 4.5) ^2^ + *z* ^2^ ≤ 2.5^2^)} and {*Ω* _*nucl*1_| (*x* ^2^ + (*y* − 12.5)^2^ + *z* ^2^ ≤ 2.5^2^)}, and the overall space *Ω* = *Ω* _*cyto*_ ⋃ *Ω* _*nucl*1_ ⋃ *Ω* _*nucl*2_ that includes the cytosol and the nuclei is defined as {*Ω* | (*x* ^2^ + (*y* − 4.5) ^2^ + *z* ^2^ ≤ 4.15^2^) ⋃ (*x* ^2^ + (*y* − 12.5) ^2^ + *z* ^2^ ≤ 4.15^2^)}.

#### Model parameters

The model approximates all processes as continuous and yields spatial distributions of Mad and pMad, both in the cytoplasm and nuclei. Rates of phosphorylation and dephosphorylation, as well as fluxes between the nuclei and cytoplasm, are assumed to be linear functions of respective concentrations, [*Mad*] and [*pMad*].

##### k_1_ and k_2_

Because enzymatic reactions occur at the membranes, their rates may also be regarded as flux densities of the corresponding volume variables, [*Mad*] and [*pMad*]. Specifically, the rate of phosphorylation is described as *k*_1_[*Mad*], where constant *k*_1_ is the product of a phosphorylation rate constant and a surface density of the activated receptor. The units of *k*_1_ are micron per second. Similarly, the rate of dephosphorylation is *k*_2_ [*pMad*], with constant *k*_2_ being the product of a dephosphorylation rate constant and the phosphatase surface density, assumed to be uniform over a nuclear envelope and same for both envelopes.

##### p^(p)Mad^ and K

The accumulation of pMad in cell nuclei is due to unequal permeabilities of a nuclear envelope for pMad import/export (Li et al., 2018; Schmierer et al., 2008). In the model, we describe the net pMad influx density as 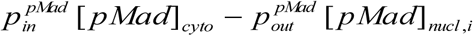, where 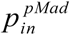 and 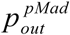 are the respective permeabilities of a nuclear envelope and index *i* denotes the nuclei of GSC (*i* = 1) and CB (*i* = 2). Consistently with a published study (Li et al., 2018), we assume 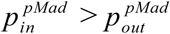. The pMad binding to the nuclear matrix and DNA is approximated by a first-order reversible reaction with the rate *k*_*on*_ ([*pMad*] − *K*[*pMad*_*bound*_]), where *k*_*on*_ is the on-rate constant in s^-1^ and *K* is the dimensionless dissociation constant. While Mad is also partially bound in the nuclei, no accumulation of Mad is observed in the nuclei of non-activated cells, likely because the effect of binding was counterbalanced by inequality of permeabilities in favor of export (Li et al., 2018; Schmierer et al., 2008). In our model, we thus ignore, for simplicity, the binding of Mad inside the nuclei and describe the net Mad influx density as *p* ^*Mad*^ ([*Mad*] _*cyto*_ −[*Mad*] _*nucl,i*_).

##### D

Diffusion of Mad and pMad molecules, both in the cytoplasm and nuclei, is described by the same diffusion coefficient *D*.

Parameter values were constrained by the data of Figure 1C, 2D, S1B and S1C, and by the estimate that pMad constitutes approximately 12.5% of total Mad; this estimate was obtained by comparing total Mad nuclear/cytoplasmic ratio between pMad positive (GSCs) and negative cells (CBs).

#### Equations

Let **r** ≡ **r**(*x, y, z*) be the radius vector of a point (*x, y, z*) in *Ω*.

The equations governing [*Mad*] and [*pMad*] in the cytosol, i.e. at the points **r** ∈*Ω*_*cyto*_, are

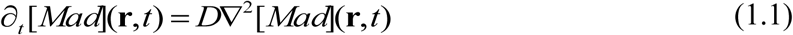

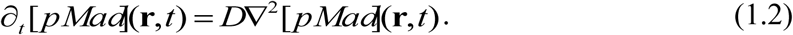

Eqs (1) are subject to boundary conditions at the plasma membrane *∂Ω* and nuclear envelopes *∂Ω*_*nucl*1_, *∂Ω*_*nucl*2_. At the plasma membrane, the boundary condition for Eq (1.1) is nonzero only for **r** ∈(*∂Ω* ⋂ *y* ≤1), the points comprising see the red ‘cap’ in Figure 3C,

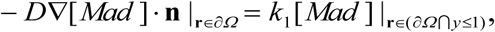

where **n** is the outward normal at *∂Ω*.

At the nuclear envelopes, the boundary conditions for [*Mad*] are

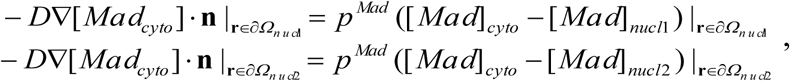

Where vectors **n** are the outward, with respect to *Ω*_*cyto*_, normals at *∂Ω*_*nucl*1_ and *∂Ω*_*nucl*2_, respectively.

Similarly, the boundary conditions for Eq (1.2) are as follows:

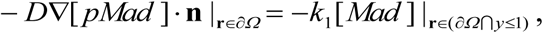

where **n** is the outward normal vector at *∂Ω*, and

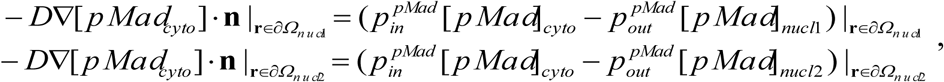

where vectors **n** are the outward, with respect to *Ω*_*cyto*_, normals to *∂Ω*_*nucl*1_ and *∂Ω*_*nucl*2_, respectively.

The equations for [*Mad*] and [*pMad*] in the nuclei, i.e. for **r** ∈(*Ω*_*nucl*1_ ⋃*Ω*_*nucl*2_), are

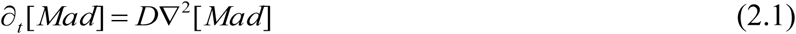

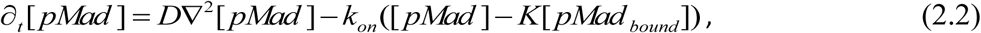

and the equation governing [*pMad bound*] is *∂*_*t*_ [*pMad*_*bound*_] = *k*_*on*_ ([*pMad*] − *K*[*pMad*_*bound*_]). Eqs (2) are subject to boundary conditions at the nuclear envelopes *∂Ω*_*nucl*1_, *∂Ω*_*nucl*2_. The boundary conditions for Eqs (2.1) are

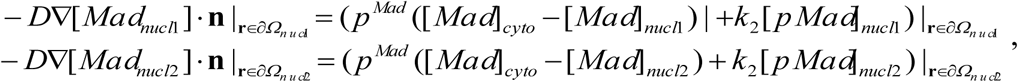

where **n** are the respective normal vectors at *∂Ω*_*nucl*1_ and *∂Ω*_*nucl*2_, outward with respect to *Ω*_*cyto*_.

Similarly, the boundary conditions for Eq (2.2) are

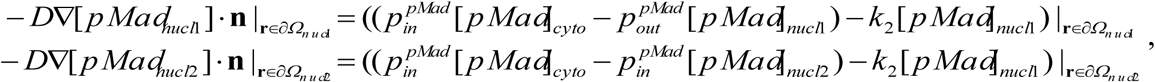

using the same definition of vectors **n** as above.

The system of Eqs (1, 2) is initialized as follows: for all **r** ∈*Ω*, [*Mad*](**r**,0) =1a. u. (an arbitrary unit of volume density) and[*pMad*](**r**,0) = 0.

The mathematical model outlined above was solved numerically with Virtual Cell (VCell), a publicly available software for computational modeling in cell biology(Slepchenko and Loew, 2010) (Resasco et al., 2012). Simulations were run with a VCell fully-implicit spatial solver on a uniform orthogonal mesh with a mesh size of 0.2 μm. The steady-state solutions were obtained by running simulations for sufficiently long end times (typically, the end time of 10^3^ seconds was sufficient to reach a steady state, with the exception of smaller values of *p* ^*Mad*^ and *k*_2_ that respectively required the end times of 2 · 10^3^ and 3 · 10^3^ seconds).

The VCell implementation of the model can be found in the VCell database of public MathModels under username ‘boris’; the model name is ‘Inaba_Model_public’. Note that in this implementation, the units of concentration are μM (1 μM = 602 molecules per μm^3^).

### Statistical analysis and graphing

No statistical methods were used to predetermine sample size. The experiments were not randomized. The investigators were not blinded to allocation during experiments and outcome assessment. Statistical analysis and graphing were performed using GraphPad prism 7 software. Data are shown as means+/-s.d. The P value (1-way ANOVA) is provided for multiple comparison with the control shown as *P<0.05, **P<0.01, ***P<0.001; NS, non-significant (P≥0.05).

### Contact for reagent and resource sharing

Further information and requests for resources and reagents should be directed to and will be fulfilled by the Contact, Mayu Inaba.

### Supplementary video

Video 1; A representative movie of Mad-GFP expressed in germline. After photobleaching of preCB side, images were taken every 10 seconds.

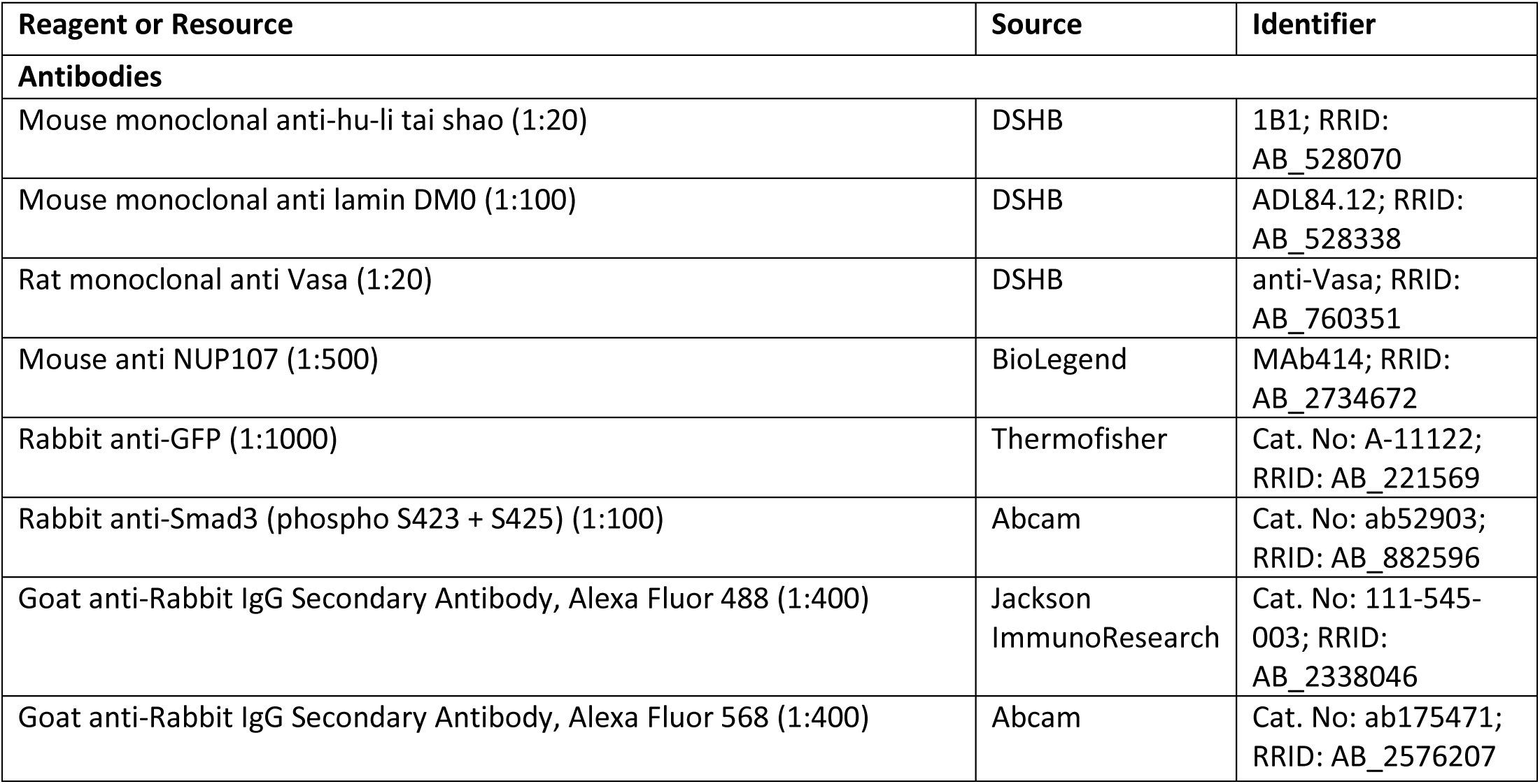
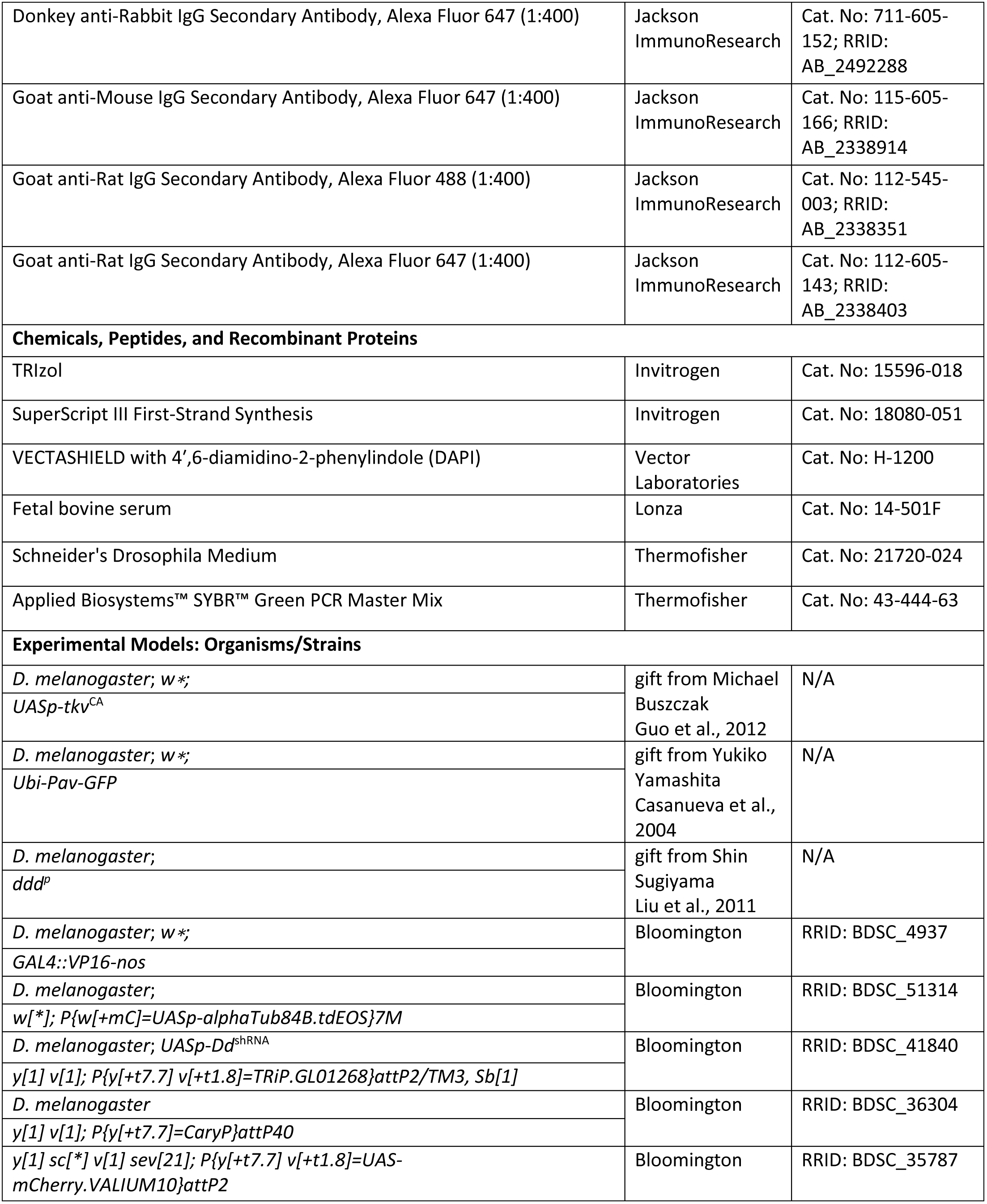
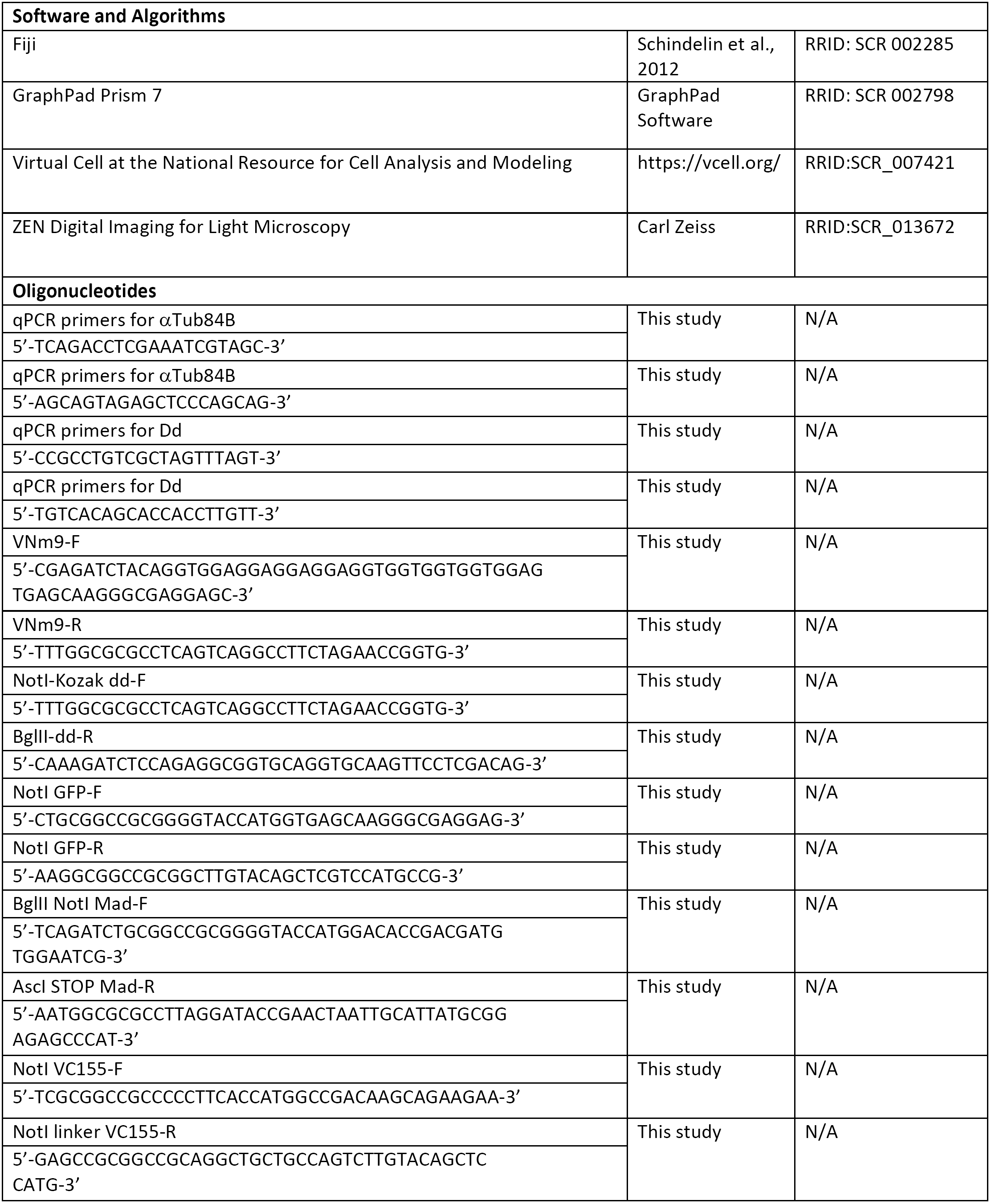
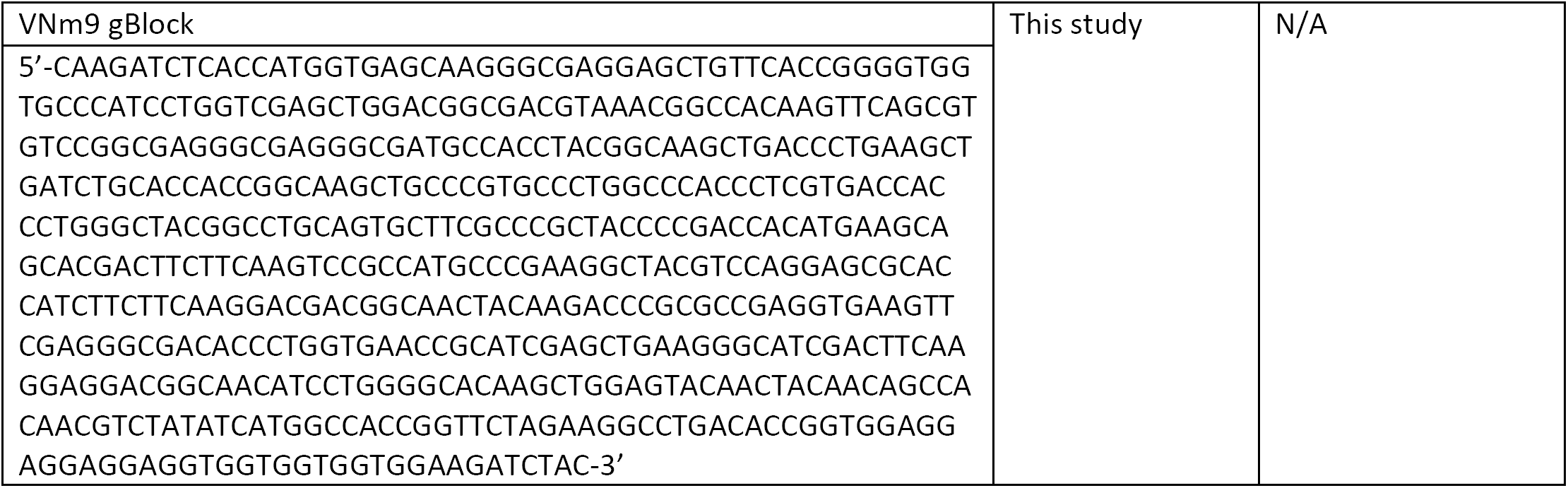

