## Supplemental Figures for "Mad dephosphorylation at the nuclear envelope is essential for asymmetric stem cell division"

### Figure S1

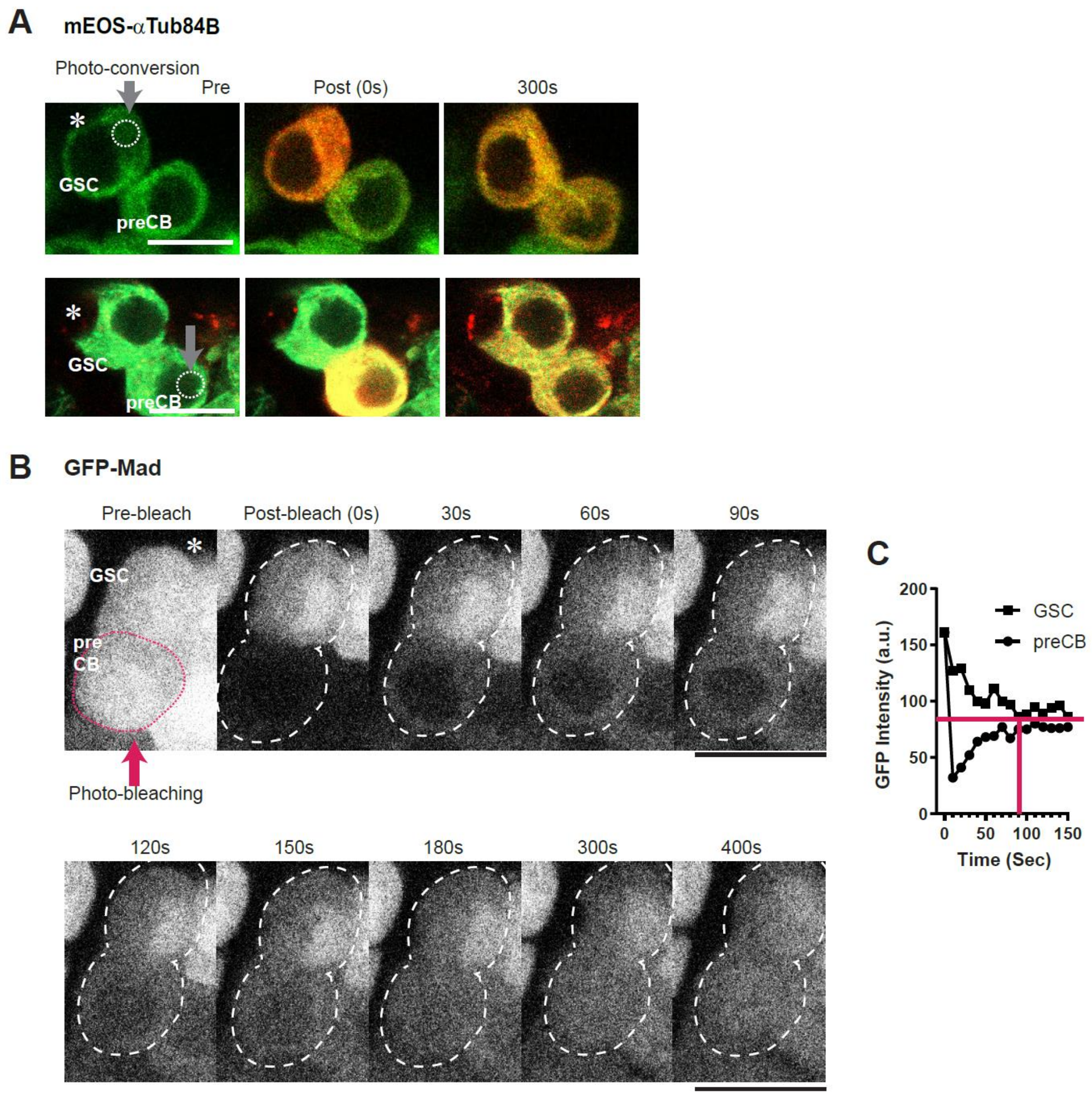

**Cytoplasmic protein diffusion between GSC and preCB**

**A,** Representative images of pre-photoconversion and after indicated recovery time (seconds) of nosGal4>mEOS-αTubulin in dividing GSC daughters. White broken circles indicate photoconverted portions. **B,** A representative image of GFP-Mad in GSC/preCB pair (encircled by white broken lines) pre-bleaching and after indicated recovery time (seconds). Pink broken lines indicate photobleached portions. **C,** The graph shows time courses of the cytoplasm intensities of GFP-Mad in GSC or preCB. Bleaching was done between time zero and 10 seconds. The pink horizontal line is manually fitted to the approximate equalized level of GFP-Mad. The vertical line indicates the approximate time when the GSC/preCB pair reached to the equal level of cytoplasmic GFP-Mad. Asterisks indicate location of CCs. Scale bars; 10 μm.

Figure S2

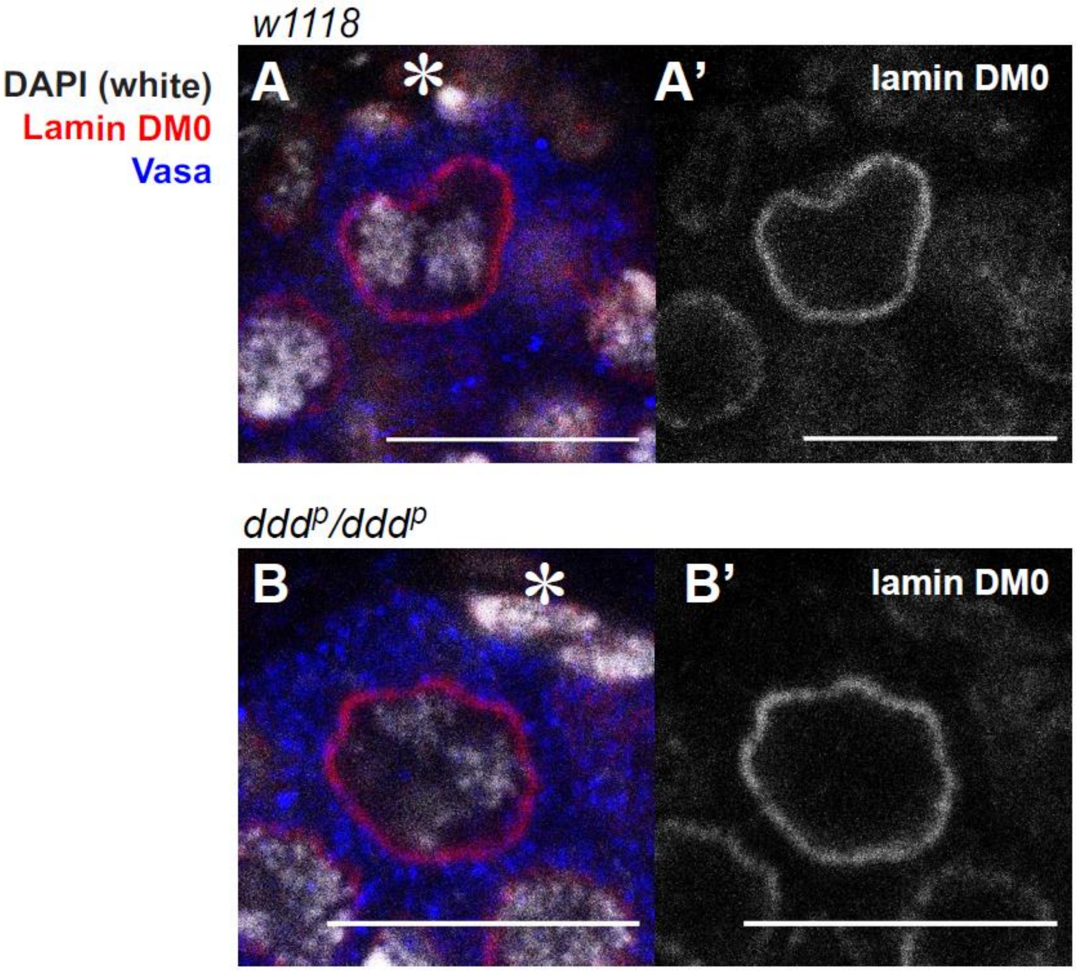


**Dd mutation does not affect nuclear envelope morphology in GSCs**

**A, B,** Representative image of Lamin DM0 (red) and Vasa staining (blue) of GSC from indicated genotypes. DAPI (white). Light panels show Lamin DM0 channel. Asterisks indicate location of CCs. Scale bars, 10 μm.

Figure S3

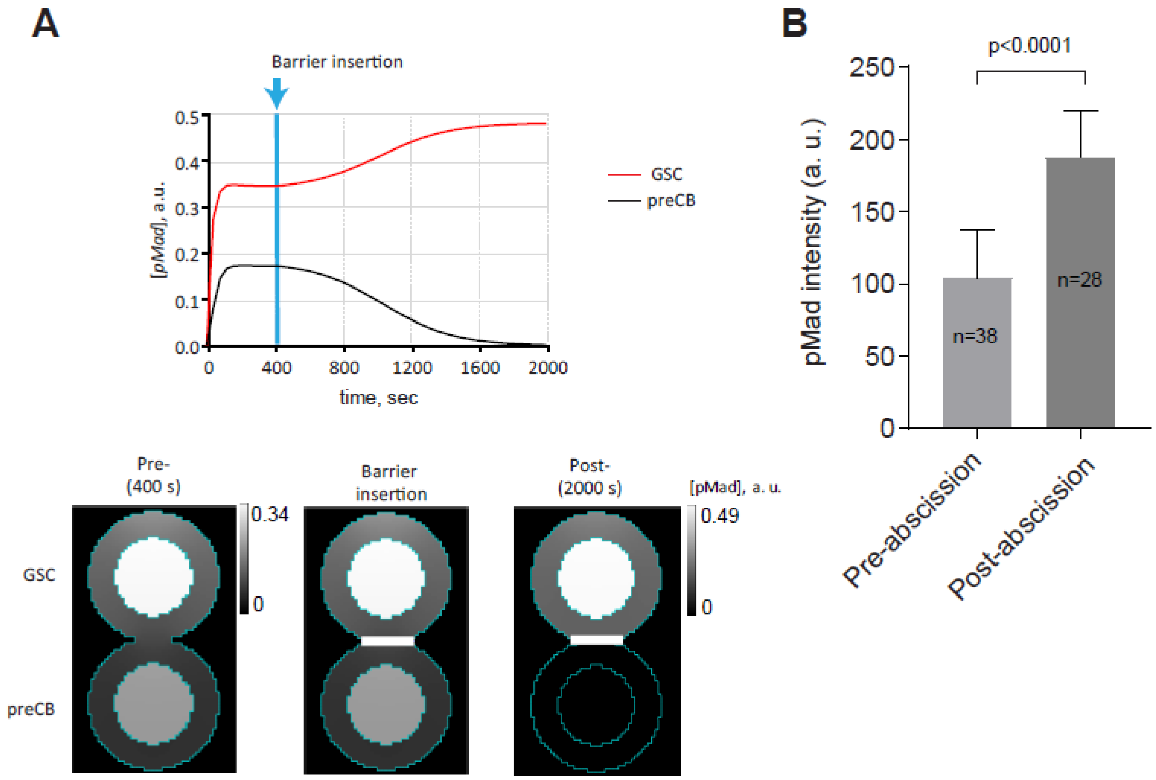


**pMad level quickly increases in GSC and decreases in preCB upon cytoplasmic separation.**

**A,** Simulated time courses of pMad concentrations in the nuclei of GSC (red) and preCB (black). Formation of a diffusion barrier started at $t$ = 400 s. The lower panel illustrates steady-state distributions of pMad before and after formation of the barrier (the white bar indicates the barrier). **B,** Comparison of pMad intensities (pMad-BG) in GSCs before and after separation from preCB.
